# Dynamic histone acetylation in floral volatile synthesis and emission in petunia flowers

**DOI:** 10.1101/2021.02.02.429379

**Authors:** Ryan M. Patrick, Xing-Qi Huang, Natalia Dudareva, Ying Li

## Abstract

Biosynthesis of secondary metabolites relies on primary metabolic pathways to provide precursors, energy, and cofactors, thus requiring coordinated regulation of primary and secondary metabolic networks. However, to date it remains largely unknown how this coordination is achieved. Using *Petunia hybrida* flowers, which emit high levels of phenylpropanoid/benzenoid volatile organic compounds (VOCs), we uncovered genome-wide dynamic deposition of histone H3 lysine 9 acetylation (H3K9ac) during anthesis as an underlying mechanism to coordinate primary and secondary metabolic networks. The observed epigenome reprogramming is accompanied by transcriptional activation, at gene loci involved in primary metabolic pathways that provide precursor phenylalanine, as well as secondary metabolic pathways to produce volatile compounds. We also observed transcriptional repression among genes involved in alternative phenylpropanoid branches that compete for metabolic precursors. We show that GNAT family histone acetyltransferase(s) (HATs) are required for the expression of genes involved in VOC biosynthesis and emission, by using chemical inhibitors of HATs, and by knocking down a specific HAT, *ELP3*, through transient RNAi. Together, our study supports that chromatin level regulatory mechanisms may play an essential role in activating primary and secondary metabolic pathways to regulate VOC synthesis in petunia flowers.

**HIGHLIGHT:** Our study shows that posttranslational modification of histones is essential for regulating the biosynthesis and emission of floral scent compounds, thus providing insights into chromatin level regulation of secondary metabolism.

## INTRODUCTION

Plants have a remarkable ability to synthesize a vast array of secondary metabolites that are not only vital for plant growth, development, reproduction, and defense, but also play crucial roles for mankind in food, medicine, industrial raw materials, and biofuels. A unique subset of secondary metabolites consists of volatile organic compounds (VOCs) that plants use for communication and interaction with the surrounding environment (Muhlemann et al., 2014a). Structurally diverse, VOCs are released from every plant tissue and play key roles in attracting pollinators and seed dispersers (Brodmann et al., 2009; Borges et al., 2010; Byers et al., 2014), above- and below-ground herbivore defense (Kessler and Baldwin, 2001; Kost and Heil, 2006), protection from pathogens and abiotic stresses (Affek and Yakir, 2002; Yi et al., 2009; Ameye et al., 2015), and plant-plant (Engelberth et al., 2004; Kost and Heil, 2006) and inter-organ signaling (Boachon et al., 2019). In most cases, emitted VOCs are comprised of complex blends of metabolites, *e.g.* there are up to 100 different compounds in floral scent bouquets (Dudareva et al., 2006). Synthesis and release of such complex mixtures requires orchestrated activation of multiple enzymatic steps among and within biosynthetic pathways.

In the last two decades, significant progress has been made in the isolation and characterization of genes responsible for the formation of plant VOCs. In general, biosynthesis of secondary metabolites relies on primary metabolic pathways, which provide precursors for their formation. It has been shown that volatiles are synthesized *de novo* in the tissues from which they are emitted and their production and emission are spatially (Dudareva et al., 1996; Kolosova et al., 2001a; Underwood et al., 2005), developmentally and/or temporally regulated (Negre et al., 2003; Colquhoun et al., 2010; Spitzer-Rimon et al., 2010; Fenske et al., 2015), and depend on biotic and abiotic factors (Qualley and Dudareva, 2008; Dudareva et al., 2013). VOC biosynthesis is mainly regulated at the level of gene expression (Dudareva et al., 1996; Gershenzon et al., 2000; Dudareva et al., 2000; Muhlemann et al., 2012), activity of enzymes responsible for the final step of VOC formation, and by substrate availability (De Moraes et al., 2001; Martin et al., 2003; Arimura et al., 2004; Effmert et al., 2005; Guterman et al., 2006; Vogel et al., 2008; Colquhoun et al., 2010; Maeda et al., 2010). Genes involved in the formation of VOCs have been shown to exhibit coordinated transcriptional activation coinciding with the VOC emission (Dudareva et al., 2003; Verdonk et al., 2005; Colquhoun et al., 2010). However, to date it has still remained largely unknown how the changes in the expression status are achieved and the regulatory mechanisms underlying this transcriptional reprogramming.

It has become increasingly clear that epigenetic regulation, including histone modification, DNA methylation, and chromatin remodeling, significantly contributes to transcriptional regulation during plant development and environmental responses (Chinnusamy and Zhu, 2009; He et al., 2011). Histone N-terminal tails are dynamically and reversibly modified with chemical groups and small proteins by histone modifying enzymes (Zhang and Reinberg, 2001). These modifications can alter the local chromatin landscape of a gene locus and are recognized by structural and regulatory proteins, which leads to transcriptional activation or silencing of the gene (Strahl and Allis, 2000; Rea et al., 2000; Musselman et al., 2012). In fungi, histone modification was shown to control gene clusters governing production of secondary metabolites (Strauss and Reyes-Dominguez, 2011; Pfannenstiel and Keller, 2019). However, our understanding of the chromatin level regulation of secondary metabolic pathways in plants remains rather limited.

*Petunia hybrida* cv. Mitchell flowers emit high levels of predominantly phenylalanine-derived phenylpropanoid/benzenoid volatiles, and subsequently have been widely used as a model system (Schuurink et al., 2006; Dudareva et al., 2013) to identify and characterize enzymes and genes involved not only in the biosynthesis and emission of scent compounds, but also in the formation of their precursor, phenylalanine (Maeda and Dudareva, 2012). In this study, we use petunia flowers to investigate chromatin level mechanisms that regulate genes involved in the formation of primary metabolite precursors and secondary metabolite volatile compounds, at a genome-wide level using *Ch*romatin *I*mmunoprecipitation *Seq*uencing (ChIP-Seq). We show that chromatin modifications during anthesis, specifically H3K9ac, facilitate the activation of the shikimate and phenylalanine synthesis pathway to provide the primary metabolite precursor, as well as distinct secondary metabolic pathways to generate the floral VOC products.

## MATERIALS AND METHODS

### Chromatin immunoprecipitation (ChIP)

Approximately two grams of corolla tissue from *P. hybrida* cv. Mitchell flowers grown under standard greenhouse conditions were harvested at the day 0 (bud) or day 2 (post-anthesis) stage at 3 p.m. in three replicates. The corolla tissue was fixed in 37 ml of 1% formaldehyde under vacuum twice, first for 15 minutes and then for 10 minutes. The reaction was stopped with addition of 2.5 ml of 2M glycine, placed under vacuum for 5 minutes, and then washed three times with ultrapure water. The fixed tissue was then dried and frozen in liquid nitrogen. Chromatin extraction and immunoprecipitation was performed as previously described (Gendrel et al., 2005) (Li et al., 2015). In detail, frozen, ground tissue was suspended in 30 ml of extraction buffer (EB1), which was filtered through Miracloth (Millipore Sigma) and then centrifuged at 4°C (1,400 *g*, 15 min). The pellet was resuspended in 1 ml of a second extraction buffer (EB2) and centrifuged again at 4°C (11,000 *g*, 15 min). The pellet was then resuspended in 300 μl of a third extraction buffer (EB3) and overlayed on 300 μl of EB3 and centrifuged at 4°C (16,000 *g*, 1 hr) to obtain the chromatin pellet. The pellet was resuspended in 300 μl of nuclei lysis buffer and sonicated with a BioRuptor Pico sonicator (Diagenode) for 26 to 36 cycles to obtain an approximately 200 bp fragment distribution. Cell debris was pelleted at 4°C (12,000 *g*, 5 min) and separated from the supernatant containing chromatin. 20 μl of the supernatant was kept as Input DNA and the rest was diluted with ChIP dilution buffer for immunoprecipitation. To perform immunoprecipitation, Protein A Dynabeads (Invitrogen) were first resuspended in ChIP dilution buffer. 50 μl of beads were used for preclearance of chromatin (rotating for 2 hr at 4°C) and during this time a second 50 μl of beads were incubated with antibody. Anti-H3K4me3 (07-473, MilliporeSigma) and anti-H3K9ac (07-352, MilliporeSigma) antibodies were used for immunoprecipitation. Precleared chromatin was then added to antibody bound beads and incubated with rotation overnight at 4°C. Beads were washed twice each with a series of buffers: low salt wash buffer, high salt wash buffer, LiCl wash buffer, and TE. Chromatin was then eluted twice with 250 μl of elution buffer at 65°C for 15 min. The two eluates were combined. The DNA crosslinking was reversed with addition of 20 μl 5M NaCl and incubation at 65°C overnight. 1 μl Proteinase K (Invitrogen) was added to digest the proteins at 45°C for 1 hr, then an equal volume of 25:24:1 phenol/chloroform/isoamyl alcohol, pH 8.0 (Invitrogen) was added to phase separate the DNA and the proteins. The DNA in aqueous phase was precipitated with sodium acetate and ethanol along with 1 μl glycogen carrier (Thermo Scientific) overnight at −20°C and then centrifuged to pellet at 4°C (12,000 *g*, 10 min). The pellet was washed with cold 70% ethanol, dried, and then resuspended in water. Concentration was measured by Qubit dsDNA High Sensitivity assay kit (Invitrogen).

### ChIP sequencing and data analysis

10 ng of input or immunoprecipitated DNA in three replicates per condition were used in library preparation for dual index Illumina paired end sequencing (Para et al., 2018) with the following modifications: (i) using NEBNext Multiplex Oligos (New England Biolabs) for adaptor ligation and barcoding primers, following manufacturer protocols; (ii) using AMPure XP beads (Beckman Coulter) for size selection; and (iii) using 18 cycles of PCR amplification with Phusion High Fidelity DNA Polymerase (New England Biolabs). Libraries were pooled and sequenced (paired-end 2×150bp) on an Illumina NovaSeq 6000 at the Purdue Genomics Core Facility. Reads were trimmed for adaptor with cutadapt (Martin, 2011) and aligned to the *Petunia axillaris* (v1.6.2) and *inflata* (v1.0.1) genomes (Fernandez-Pozo et al., 2015a; Bombarely et al., 2016) with Bowtie 2 (Langmead and Salzberg, 2012). Properly paired read alignments were then converted to bed format representing each insert fragment using bedtools (Quinlan and Hall, 2010). The aligned fragments were then used to call differential enrichment of sequencing peaks for H3K4me3 and H3K9ac ChIP between day 2 and day 0 using corresponding input DNA library as a background. For peak calling, SICER-df.sh (Zang et al., 2009; Xu et al., 2014) was run for all *Petunia* scaffolds containing at least one annotated gene, with a gap size of 200, a window size of 200, an effective genome size of 0.9, and with an FDR < 0.01 cutoff for significance. Significantly increased or decreased peaks at a two-fold or higher were intersected with annotated Petunia gene features using bedtools (Quinlan and Hall, 2010). Genes overlapped with significant peaks in all three replicates were considered significant differentially modified genes (DMGs). For downstream analyses of gene set enrichment, *P. axillaris* gene features were used as a background. The significance of overlapping gene sets between ChIP-Seq and RNA-Seq analyses were evaluated by hypergeometric test using phyper function in R. Integrative Genomics Viewer (IGV) was used for visualization of ChIP-Seq data (Robinson et al., 2011).

### RNA sequencing analysis

RNA sequencing data for day 0 and day 2 petunia corolla tissue was obtained from NBCI Gene Expression Omnibus [GSE70948] (Widhalm et al., 2015). Reads were trimmed with an in-house python script and aligned to the *Petunia axillaris* (v1.6.2) and *inflata* (v1.0.1) genomes (Fernandez-Pozo et al., 2015a; Bombarely et al., 2016) using TopHat2 (Kim et al., 2013). Aligned reads were assigned to petunia genes with HTSeq (Anders et al., 2015). Differentially expressed genes were determined using DESeq2 (Love et al., 2014) with cutoffs of |log2 fold-change| > 1 and false discovery rate (FDR) < 0.05.

### Gene Ontology enrichment analysis

In order to determine enriched Gene Ontology (GO) terms in gene sets of interest in petunia, the best *Arabidopsis thaliana* homolog to each *P. axillaris* protein was determined using blastp against the Araport11 protein database (Cheng et al., 2017) with an E-value cutoff of 1.0e-08 by the BLAST+ toolkit (Camacho et al., 2009). GO term annotations for *A. thaliana* genes curated by TAIR (Berardini et al., 2004) were then assigned to the *P. axillaris* homologs to populate a GO term background for petunia. The significant enrichment of GO terms in gene sets of interest versus the background were detected using an in-house python script, where a hypergeometric test (SciPy stats library, https://www.scipy.org) was used to calculate statistical significance of enrichment, and FDR correction (Benjamini and Hochberg, 1995) was applied to control for multi-testing error using the python module ‘statsmodels’ (Seabold and Perktold, 2010). REVIGO was used for visualization of enriched GO terms (Supek et al., 2011). Enrichment levels of statistically significant GO terms were used to create heatmaps with the R packages ‘pheatmap’ (https://cran.r-project.org/web/packages/pheatmap/index.html) and ‘RColorBrewer’ (https://cran.r-project.org/web/packages/RColorBrewer/index.html).

### Metabolic pathway analysis

Aromatic amino acid biosynthesis and phenylpropanoid pathway genes in *P. axillaris* were identified based on supplementary notes to the publication of petunia genomes (Bombarely et al., 2016) or collected knowledge of the pathways (Zhao and Dixon, 2011; Maeda and Dudareva, 2012; Dudareva et al., 2013; Muhlemann et al., 2014b; Passeri et al., 2016) with members of each pathway identified by BLAST homology. Two putative *CAD* genes were identified in *P. axillaris* based on homology to the major *CAD* genes contributing to lignin biosynthesis in *A. thaliana* (Sibout et al., 2005) and *Nicotiana attenuata* (Kaur et al., 2012). The petunia CAD1 and CAD2 proteins have 92% and 78% identity with the *N. attenuata* CAD1 protein, respectively. Putative peroxidase and laccase genes involved in lignin polymerization were identified by BLAST homology to genes identified in *A. thaliana* (Zhao et al., 2013). Significance of overlaps between metabolic pathway genes and gene lists from ChIP-Seq and RNA-Seq analyses was evaluated by hypergeometric test in R. Overlaps between gene sets were visualized using Circos (Krzywinski et al., 2009).

### Chemical inhibition assays

For histone acetyltransferase inhibitor assays, *P. hybrida* cv. Mitchell flowers grown under standard greenhouse conditions were harvested at the day 0 (bud) stage at 3 p.m. and floated in a 3% sucrose solution with mock treatment (0.1% DMSO) or histone acetyltransferase inhibitor C646 (MilliporeSigma) or MB-3 (MilliporeSigma) at a final concentration of 100 μM in 0.1% DMSO. Treatment with inhibitors did not affect anthesis, as all flowers were open by afternoon of Day 1. Corolla tissue was harvested and frozen in liquid nitrogen one hour before end of light period at 7 p.m. in 4 biological replicates of two flowers each, at day 0, day 1, and day 2. Gibberellic acid inhibitor assays were performed similarly, with uniconazole (Lkt Laboratories) or flurprimidol (bioWORLD) at a final concentration of 100 μM in 0.1% DMSO, with a single end point harvest at day 2. RNA was extracted with RNeasy Plant Mini Kit (Qiagen), treated with recombinant, RNase-free DNase I (Roche) at 37°C for one hour, followed by a cleanup with RNeasy Mini Kit (Qiagen). SuperScript IV (Invitrogen) was used to generate cDNA with oligoDT primer. qRT-PCR was performed with PowerUp SYBR Green (Applied Biosystems) using primers designed to specification with Primer3 (Untergasser et al., 2012, 3) with sequences listed (Supplementary Table S1) or as previously described for *UBQ10* and *PhABCG1* (Adebesin et al., 2017). Relative gene expression was normalized to two reference genes, *FBP1* and *UBQ10*. For ChIP-qPCR to confirm chemical inhibition of histone acetyltransferase activity, flowers were treated with mock or MB-3 as above and at day 2 were used for ChIP as described above for ChIP-Seq, with anti-H3K9ac (07-352, Millipore Sigma) and anti-H3K14ac (07-353, Millipore Sigma) antibodies used for immunoprecipitation. Primers designed to gene body acetylation peaks (Supplementary Table S1) were used for qPCR with PowerUp SYBR Green (Applied Biosystems) in order to determine pulldown percent relative to input, with normalization to housekeeping genes *ACTIN* (Peaxi162Scf00583g00529) and *PP2AA3* (Peaxi162Scf00382g00122).

### ELP3 down-regulation through transient RNAi

A 300 bp sequence specific for RNA interference mediated silencing of ELP3 was chosen using the Sol Genomics Network VIGS (Virus-Induced Gene Silencing) tool (Fernandez-Pozo et al., 2015b) and then amplified from *P. hybrida* cDNA using primers including an *attB* recombination site. The fragment was then cloned into the pDONR/Zeo gateway vector (Life Technologies) following manufacturer instructions and subcloned into the destination vector pK7GWIWG2(II) in a sense and antisense direction to facilitate hairpin formation by Invitrogen Gateway LR Clonase II Enzyme mix (Life Technologies). Binary vectors were transformed into Agrobacterium tumefaciens strain GV3101 with freeze-thaw cycles. Transient *ELP3* down regulation was achieved similarly as previously described (Yoo et al., 2013) using vacuum infiltration of at least 18 petunia flowers at day 1 stage, with transformed Agrobacterium containing pK7GWIWG2(II)-ELP3 or pK7GWIWG2(II) empty vector control at OD_600_ of 0.8. Infiltrated flowers were kept in 5% sucrose solution in dark for an additional 48 hours before floral scent was collected for 4◻hours from 17:00 to 21:00. Following scent collection, total RNA was extracted from petunia flowers using the Spectrum Plant Total RNA Kit (Sigma-Aldrich). About one microgram of total RNA was reverse transcribed to first strand cDNA in 10 μl reaction using the EasyScript cDNA synthesis kit (Applied Biological Materials) and used for qRT-PCR as described above.

### Targeted metabolite profiling

Petunia volatiles were collected from infiltrated flowers by a closed-loop stripping method and analyzed by GC-MS (Widhalm et al., 2015). Briefly, floral scents were absorbed in VOC collection traps containing 20 mg of Porapak Q (80-100 mesh) matrix (Waters). VOCs were eluted from scent collection traps with 200 μl dichloromethane. One microgram of naphthalene was added to the eluted sample as an internal standard. Samples were analyzed on an Agilent 6890N-5975B GC-MS system. Quantitation of different volatile compounds was performed based on standard curves generated with commercially available standards.

## RESULTS

### Coordinated chromatin modification and transcriptomic reprogramming in petunia corolla during flower development

In petunia flowers, floral VOCs are produced primarily in the corolla. The levels of phenylalanine precursor and VOCs are developmentally co-regulated during the lifespan of the flower, achieving a maximum on the second day post-anthesis (Verdonk et al., 2005; Maeda et al., 2010). We hypothesized that a genome-wide reprogramming of histone modifications occurs in the corolla during floral development to promote active transcription of genes involved in VOC synthesis, including the primary and secondary metabolite pathways. To test this hypothesis, corolla tissues of *Petunia hybrida* cv. Mitchell were collected for ChIP-Seq in three biological replicates, at two time points representing the lowest and highest VOC production: immediately pre-anthesis buds (day 0) and open flowers (day 2). As activation of VOC pathway genes was particularly of interest, histone marks associated with transcriptional activation, H3K9ac and H3K4me3, were chosen as the focus of this study (Ha et al., 2011; Du et al., 2013). The ChIP-Seq reads were generated using Illumina NovaSeq platform and then mapped to parental genomes of *P. hybrida*, *P. axillaris* and *P. inflata* (Bombarely et al., 2016). The majority reads mapped to the *P. axillaris* genome (Supplementary Figure S1), which agrees with previous observations (Bombarely et al., 2016). Therefore, *P. axillaris* gene features were used for downstream bioinformatic analysis to provide an appropriate statistical background. Genes associated with significantly increased or decreased levels of H3K9ac or H3K4me3 at day 2 relative to day 0 in all three biological replicates were determined using SICER (FDR < 0.01, fold change > 2) (Zang et al., 2009) and referred to as differentially modified genes (DMGs) hereinafter.

Overall, we identified a widespread reprogramming of histone modifications in petunia corolla from the bud stage (day 0) to open flower (day 2), in the form of increased levels of H3K9ac or H3K4me3 at hundreds to thousands of gene loci. Specifically, 4,207 DMGs exhibit increased H3K9ac from day 0 to day 2, and 355 DMGs displayed decreased H3K9ac (Supplementary Dataset S1). For H3K4me3, we identified 855 DMGs with increased H3K4me3 and nine DMGs with reduced H3K4me3 (Supplementary Dataset S1). To determine whether specific biological processes are associated with these DMGs, we developed an in-house Gene Ontology (GO) enrichment analysis pipeline for petunia and identified GO terms significantly enriched (FDR < 0.05) among DMGs (Figures 1A and 1B; Supplementary Tables S2-3). Interestingly, DMGs with increased H3K9ac were uniquely enriched with ‘aromatic amino acid biosynthesis’ (Figure 1A, Supplementary Table S2), suggesting that H3K9ac, but not H3K4me3, is specifically involved in activating phenylalanine biosynthesis to provide precursor for floral VOCs. Indeed, we observed a significant increase of H3K9ac levels from day 0 to day 2 at gene loci involved in the shikimate pathway and general phenylpropanoid pathway that synthesize the precursor metabolites phenylalanine and phenylalanine derivatives for VOC formation (Figure 1C: *EPSPS1, DAHP1, PALa, 4CLa*). Furthermore, we observed increased H3K9ac at gene loci involved in the formation of VOCs including volatile benzenoid/phenylpropanoid (Figure 1C; *CNL*) and volatile phenylpropene (Figure 1C; *HCT*). In addition, DMGs with increased H3K9ac are enriched with GO terms related to post-transcriptional regulation, hormone signaling, and intracellular protein transport, while DMGs with increased H3K4me3 are enriched with GO terms for post-transcriptional regulation and transport, and tissue development and cell cycle (Figure 1A&B, Supplementary Table S2-3). There was no significant GO term enriched among genes showing decreased levels of H3K9ac or H3K4me3.

**Figure 1.**
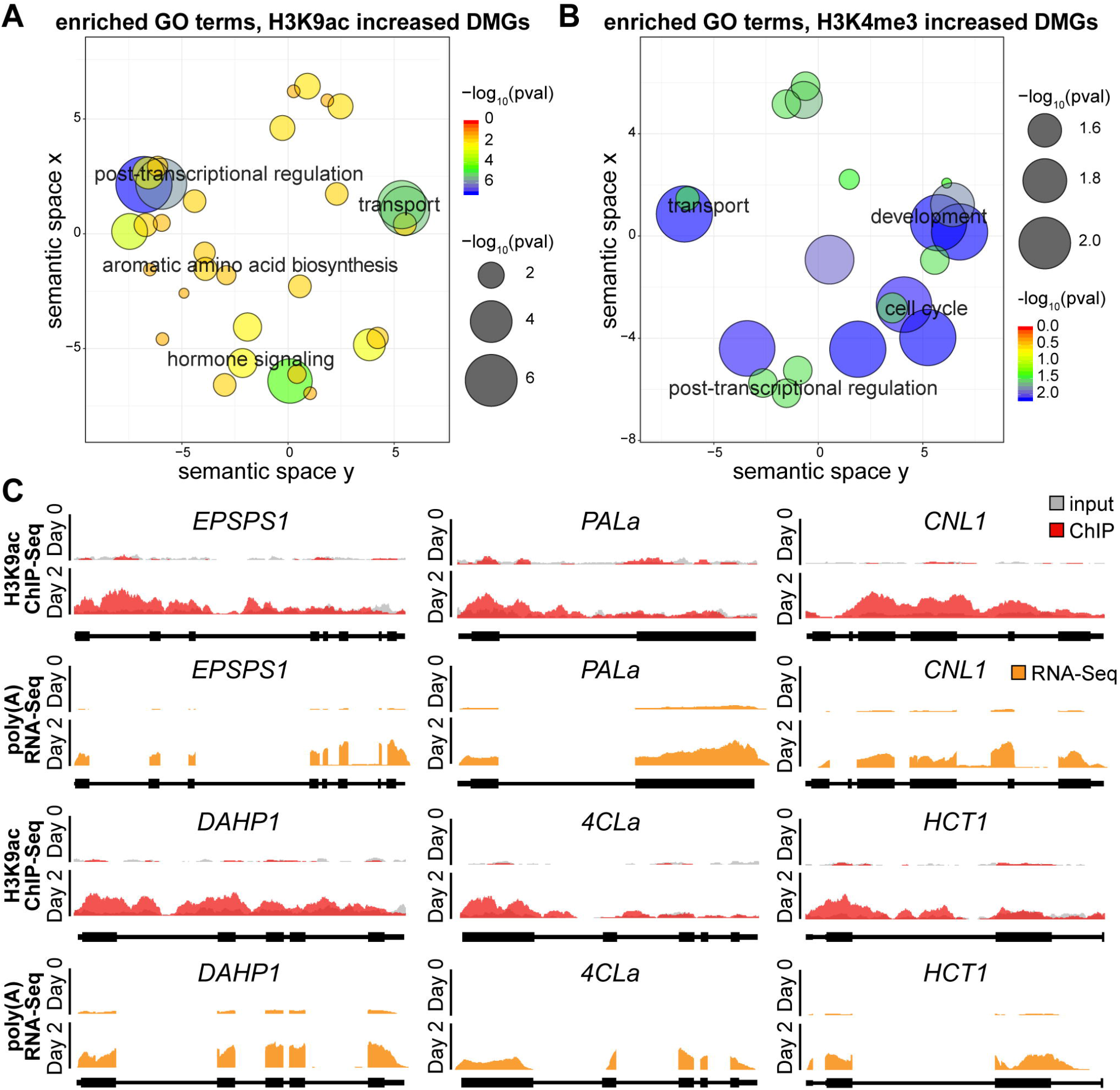
Dynamic regulation of histone modifications during anthesis in petunia corolla. Significant GO terms enriched among DMGs with increased H3K9ac (A) or increased H3K4me3 (B) on day 2 vs day O in petunia corolla are shown in sematic spaces generated by REVIGO. Each circle in the sematic space represents a significant GO term, and the size and color of the circle represent the level of significance of the enrichment. (C) Histone ChIP-Seq and RNA-Seq coverage showing dynamic H3K9ac deposition along gene body of specific VOC genes from day O to day 2 and coincident increase in mRNA level. The sequencing depth was scaled to library size. Representative results from one of three independent biological replicates are shown. 4CL, 4-coumaryl-CoA ligase; CNL, cinnamoyl-CoA ligase; DAHP, 3-deoxy-D-arabino-heptulosonate 7-phosphate synthase; EPSPS, 5-enolpyruvylshikimate 3-phosphate synthase; HCT, hydroxycinnamoyl-CoA:shikimate/quinate hydroxycinnamoyl transferase; PAL, phenylalanine ammonia lyase.

To understand the effects of observed histone modification patterns on gene expression, RNA-Seq data generated from the same tissue and same developmental stages as the ChIP-Seq were analyzed (Widhalm et al., 2015). We found a global reprogramming of gene expression in the corolla from day 0 to day 2, including 3,662 differentially expressed genes (DEGs) that are up-regulated and 4,251 that are down-regulated, determined by DESeq2 with a cutoff of FDR < 0.05 and fold change > 2 (Love et al., 2014) (Supplementary Dataset S2; Supplementary Figure S2). As expected, the six representative Phe and VOC biosynthetic genes in Fig. 1C also showed significantly increased mRNA levels (Fig. 1C), suggesting that the *hyper-*acetylation of H3K9ac is associated with increased expression of these genes. Indeed, aromatic amino acid biosynthesis was enriched among up-regulated DEGs, while flavonoid and anthocyanin biosynthesis were enriched among down-regulated DEGs; interestingly, both up- and down-regulated DEGs were significantly enriched with genes involved in cell wall modification (Figures 2A and 2B, Supplementary Table S4; will be explained in detail later). In addition, up-regulated DEGs were enriched with genes involved in post-transcriptional regulation, stress response, and hormone signaling (Figure 2A, Supplementary Table S4), while down-regulated DEGs were enriched with light response (Figure 2B, Supplementary Table S4).

**Figure 2.**
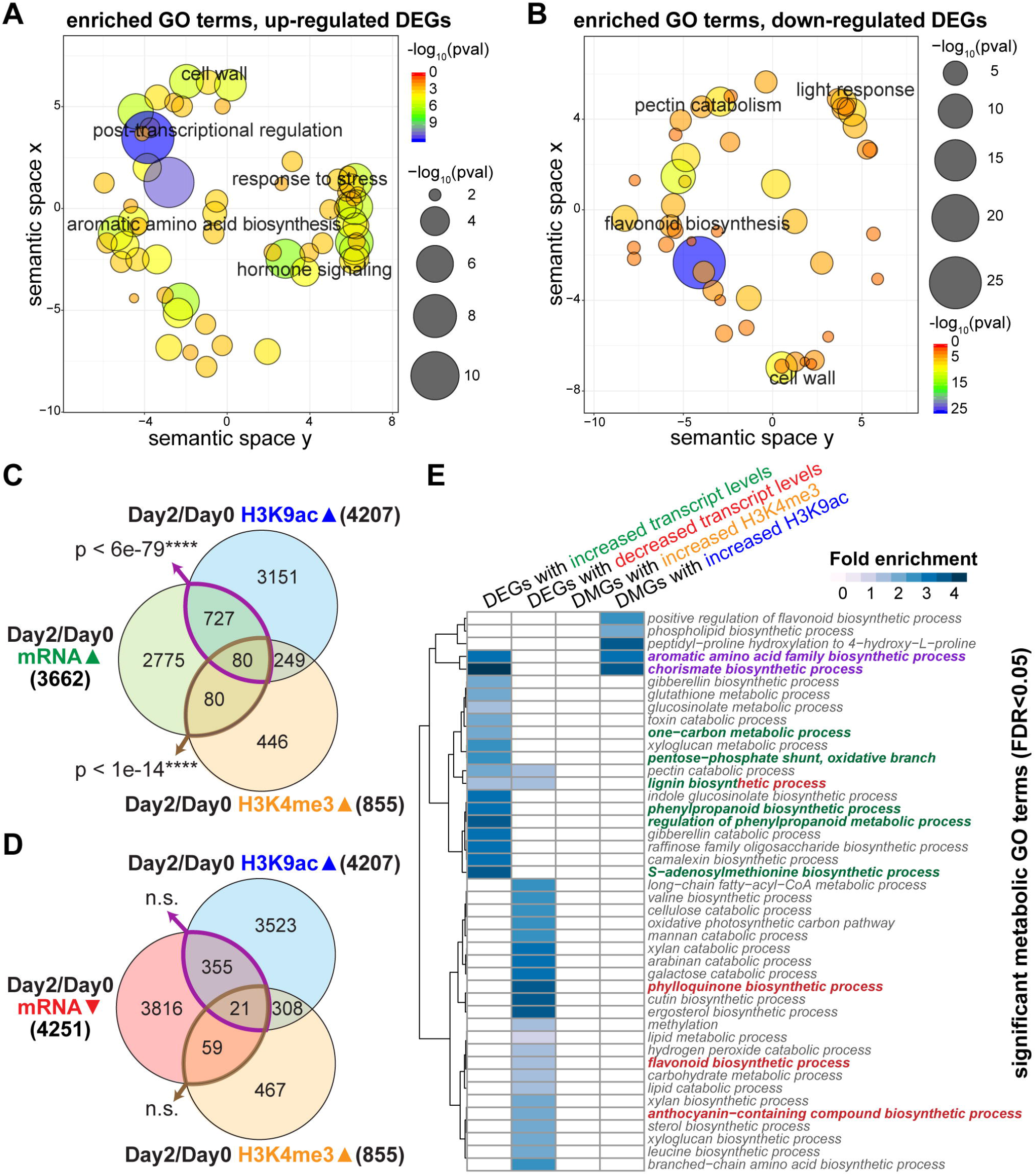
Transcriptionally and epigenetically regulated genes and metabolic pathways during anthesis in petunia corolla. Significant GO terms enriched among DEGs with increased transcript levels (A) or decreased transcript levels (B) on day 2 vs day O in petunia corolla are shown in sematic spaces generated by REVIGO. Each circle in the sematic space represents a significant GO term, and the size and color ofthe circle represent the level ofsignificance ofthe enrichment. Overlaps between sets ofDMGs with increased H3K9ac or H3K4me3 levels and DEGs with increased (C) or decreased (D) transcript levels were shown in venn diagrams. P value for the intersection between gene sets was determined by hypergeometric test. **(E)** Heatmap depicting the fold enrichment ofmetabolic GO terms significantly enriched among the DEGs or DMGs. White color represents a lack ofsignificant enrichment. Phenylpropanoid and VOC synthesis related metabolic terms are highlighted in green for up-regulated processes, red for down-regulated processes, and purple for biological processes regulated at both transcript levels and chromatin levels.

To investigate the extent to which changes of histone modifications at a gene locus co-occur with changes in transcript level, the DMGs identified by ChIP-Seq were compared to DEGs determined by RNA-Seq. Significant overlaps were observed between DEGs with increased transcript levels and DMGs that gained either H3K9ac or H3K4me3 on day 2 *vs* day 0 (Figure 2C). By contrast, genes that gain H3K9ac or H3K4me3 have no significant overlap with down-regulated DEGs (Figure 2D). Genes associated with reduced H3K9ac did not show significant overlap with either set of DEGs (Supplementary Figure S3). Overall, our results indicate that increased H3K9ac and H3K4me3 are associated with gene activation during anthesis in petunia flowers.

### Transcriptional and epigenetic regulation of phenylpropanoid metabolism in petunia flowers

To further investigate the regulation of biosynthesis and emission of floral VOCs, we first examined in detail the enriched metabolic GO terms among DEGs and DMGs for their relevance to floral VOC emission (Figure 2E). Interestingly, only two significantly over-represented GO terms were found to be shared between DMGs (with increased H3K9ac) and DEGs (up-regulated): ‘chorismate biosynthetic process’ and ‘aromatic amino acid family metabolic process’ (Figure 2E, GO terms in purple). Increased flux through the shikimate pathway to promote production of chorismate and subsequently phenylalanine, the predominant precursor of petunia floral VOCs, is known to be an important factor contributing toward VOC production (Colquhoun et al., 2010; Oliva et al., 2015; Schuurink et al., 2006; Widhalm at al., 2015). Our integrated analysis now revealed that this crucial process is regulated at both epigenetic and transcript levels.

In addition to the production of precursors chorismate and phenylalanine, more significant GO terms relevant to floral VOC synthesis were identified among the up-regulated DEGs, including ‘phenylpropanoid biosynthetic process’ and ‘regulation of phenylpropanoid metabolic process’ (highlighted in green in Figure 2E). The term ‘pentose-phosphate shunt, oxidative branch’ is also of relevance, as the pentose phosphate pathway provides erythrose 4-phosphate (E4P) precursor for the shikimate pathway to produce chorismate and subsequently phenylalanine (Maeda and Dudareva, 2012). Also present as enriched GO terms are ‘S-adenosylmethionine (SAM) biosynthetic process’ and the connected ‘one-carbon metabolic process’. SAM is an methyl donor in the production of volatile metabolites such as methylbenzoate and eugenol (Kolosova et al., 2001b; Shaipulah et al., 2016). It has been shown that *SAM SYNTHETASE* is regulated coordinately with VOC and shikimate pathway enzymes in petunia during flower development (Verdonk et al., 2003; Verdonk et al., 2005). Here, our transcriptome analysis expanded the previous reports to genome-wide level, revealing that multiple genes in the SAM biosynthetic pathway are targets of transcriptional activation during corolla maturation and VOC synthesis.

GO term analysis of down-regulated DEGs revealed transcriptional repression of metabolic pathways that compete with VOC metabolism for precursors or cofactors, including ‘flavonoid biosynthetic process’, ‘anthocyanin-containing compound biosynthetic process’, and ‘phylloquinone biosynthetic process’ (Figure 2E, terms highlighted in red). Phylloquinone is produced from chorismate, while flavonoids and anthocyanins are derivatives of phenylalanine and share metabolic intermediates with VOC pathways (Maeda and Dudareva, 2012; Bombarely et al., 2016). Production of high levels of VOCs in *P. hybrida* flowers is therefore coincident with the reduced expression of genes involved in competing pathways that consume phenylpropanoid metabolic precursors. Interestingly, the GO term for ‘lignin biosynthetic process’ is present among up-regulated DEGs as well as down-regulated DEGs (Figure 2E).

Overall, our analysis of GO terms revealed coordinated H3K9ac deposition at gene loci involved in the chorismate and aromatic amino acid metabolic pathways in petunia flowers, accompanied by transcriptional activation of these pathways. We also showed that global transcriptional reprogramming deactivates phenylpropanoid metabolic branches that would draw phenylalanine or other phenylpropanoid intermediates away from VOC production such as anthocyanin/flavonoid biosynthesis pathways. However, these enrichment analyses are based on GO annotations propagated from the Arabidopsis model, thus lineage-specific secondary metabolism genes involved in floral VOC synthesis and emission are overlooked. Moreover, we were interested in identifying the specific points of transcriptional and chromatin regulation within primary and secondary metabolic networks. Therefore, we further examined in detail individual metabolic branches and pathways related to VOC metabolism for transcriptional and epigenetic regulatory targets, based on an extensive collection of curated knowledge of these pathways in petunia (Supplementary Tables S5-7), detailed as below.

### Shikimate and phenylalanine biosynthesis pathways

The enzymes involved in the shikimate and phenylalanine biosynthesis pathways have been well characterized in plants (Figure 3A, Supplementary Table S5) (Maeda et al., 2010; Maeda et al., 2011; Maeda and Dudareva, 2012; Yoo et al., 2013; Qian et al., 2019). Overlaying the epigenomic and transcriptomic data revealed systematic H3K9ac deposition and transcriptional activation at almost every step of the pathway, for at least one gene copy if multiple genes are potentially involved in driving an enzymatic step (Figure 3A). In total, 46% of the genes in this pathway display increased H3K9ac, and of those 73% show increased gene expression (Figure 3B). Statistically, genes encoding enzymes in this pathway are significantly enriched with DMGs gaining H3K9ac (overlap *p*-value < 3.34E-05) and DEGs with increased transcript levels (overlap *p*-value < 9.67E-09); by contrast, genes with change in H3K4me3 levels do not appear significantly enriched. Our results therefore support that flux through the shikimate and phenylalanine biosynthetic pathways is a major target of H3K9ac and transcriptional regulation during petunia flower development to potentiate VOC production.

**Figure 3.**
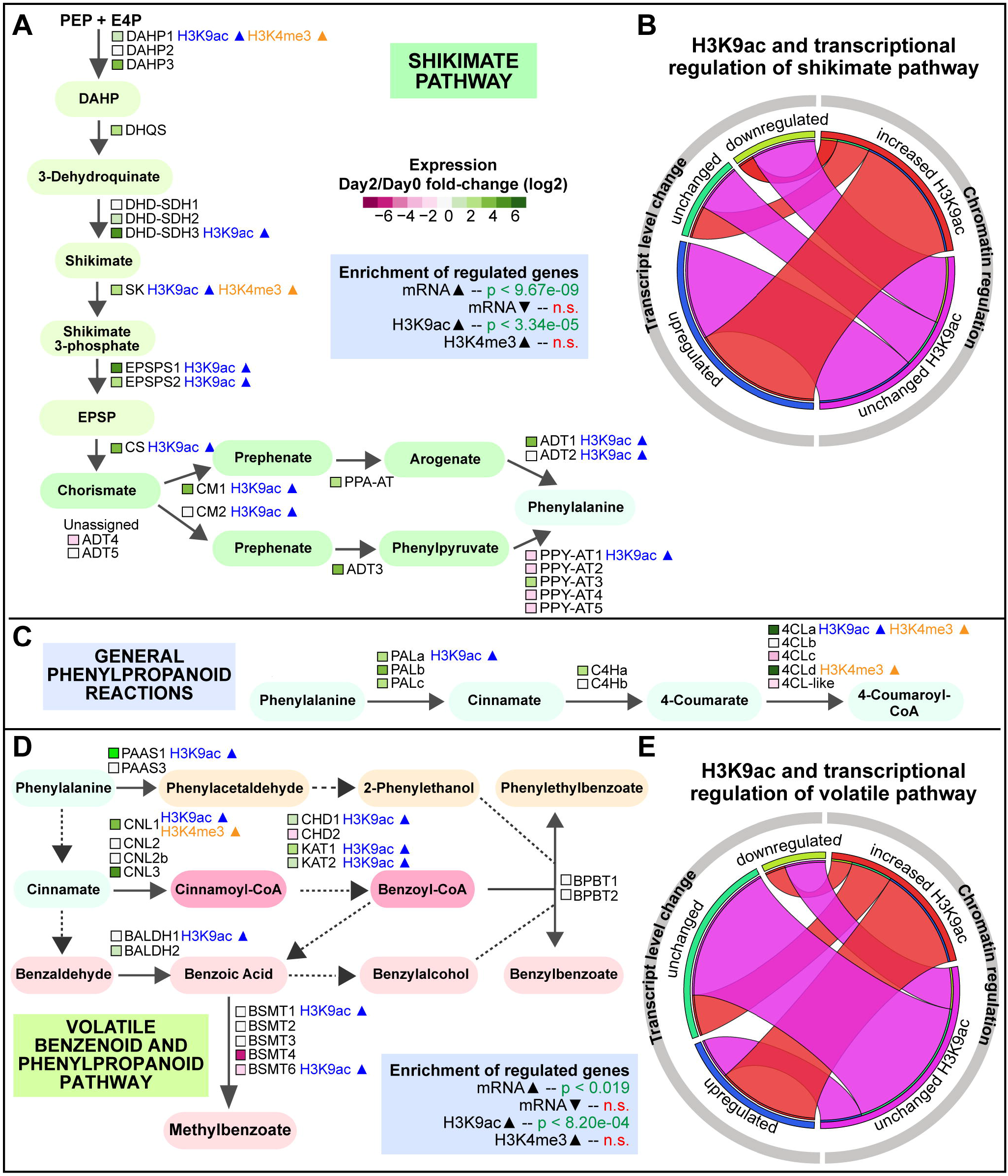
Transcriptional and epigenetic regulation of the shikimate, general phenylpropanoid, and volatile benzenoid/phenylpropanoid pathways. **(A)** Schematic of the shikimate and phenylalanine biosynthesis pathway with transcriptomic and epigenomic data overlaid. Color legend: shikimate pathway intermediates in light green, chorismate and phenylalanine synthesis intermediates in blue-green, and phenylalanine in light blue. The transcript level changes of genes underlying each enzymatic step are represented by the colored squares with color key shown in the graph (while green represents increased transcript levels and magenta represents decreased transcript levels). The significance of overlaps with DEGs and DMGs are determined by hypergeometric tests and are shown in the blue shaded box. **(B)** Circos plot showing the two regulatory categories (transcriptional level changes and chromatin regulation) for genes involved in the shikimate pathway, as well as intercategorical relationships between the two regulatory categories. The plot consists of two semicircles. The semicircle on the left represents the transcriptional regulation categories for genes involved in the shikimate pathway: the number of genes that are down-regulated, up-regulated, or unchanged are proportional to the length of the three curved segments in dark blue, turquoise, and green, respectively. Similarly, the semicircle on the right represents two chromatin regulatory categories for H3K9ac (increased, or unchanged), for the same set of genes involved in shikimate pathway. The genes shared between a transcriptional regulatory category (e.g. up-regulated) and a chromatin regulatory category (e.g. increased H3K9ac) are represented by the ribbon linking the two curved segments located within the two semicircles. For example, majority of the genes with increased H3K9ac are also up-regulated at transcript levels. **(C)** Schematic of the general phenylpropanoid genes generating shared metabolic intermediates (in light blue) with transcriptomic and epigenomic data overlaid. **(D)** Schematic of the volatile benzenoid and phenylpropanoid biosynthesis pathway with transcriptomic and epigenomic data overlaid for known genes. Color legend: general phenylpropanoid intermediates phenylalanine and cinnamate in light blue, phenylpropanoid-related (C6-C2) derived volatiles in orange, phenylpropanoid (C6-C3) and benzenoid (C6-Cl) intermediates in magenta and derived volatiles in pink. **(E)** Circos plot showing the two regulatory categories (transcriptional level changes and H3K9ac changes) and intercategorical relations of volatile benzenoid and phenylpropanoid pathway genes. 4CL, 4-coumaryl-CoA ligase; ADT, arogenate dehydratase; BALDH, benzaldehyde dehydrogenase; BP B T, benzoyl-CoA:benzylalcohol/phenylethanol benzoyltransferase; BSMT, benzoic acid/salicylic acid carboxyl methyltransferase; C4H, cinnamate 4-hydroxylase; CHD, cinnamoyl-CoA hydratase-dehydrogenase; CM, chorismate mutase; CNL, cinnamoyl-CoA ligase; CS, chorismate synthase; DAHP, 3-deoxy-D-arabino-heptulosonate 7-phosphate; EPSPS, 5-enolpyruvylshikimate 3-phosphate; DHD-SDH, 3-dehydroquinate dehydratase and shikimate dehydrogenase; DHQS, 3-dehydroquinate synthase; KAT, 3-ketoacyl-CoA thiolase; PAAS, phenylacetaldehyde synthase; PAL, phenylalanine ammonia lyase; PPA-AT, prephenate aminotransferase; PPY-AT, phenylpyruvate aminotransferase; SK, shikimate kinase.

### General phenylpropanoid reactions

From phenylalanine, a set of reactions catalyzed by PAL, C4H and 4CL generates metabolites shared by many downstream phenylpropanoid branches, including those toward synthesis of flavonoids, lignin, and VOCs (Ferrer et al., 2008; Bombarely et al., 2016) (Figure 3C; Supplementary Table S5). In petunia, these enzymes are encoded by multiple gene loci (Bombarely et al., 2016), with many showing increased transcript levels on day 2 relative to day 0 (Figure 3C). H3K9ac deposition during anthesis is observed at one copy of *PAL* and one copy of *4CL*. In the case of *4CL*, two of the five copies are transcriptionally increased. Interestingly, the two induced copies also display increased H3K9ac or H3K4me3. 4CL drives the synthesis of 4-coumaryl-CoA, which has multiple fates in metabolism and can serve as a precursor for flavonoids and anthocyanins, lignin, and phenylpropene VOCs. The distinct chromatin and transcriptional regulation of these five copies could possibly indicate specialized gene functions for different metabolic fates.

### Volatile benzenoid and phenylpropanoid pathways

The majority of volatile phenylpropanoid (C_6_-C_3_) and benzenoid (C_6_-C_1_) compounds are synthesized from cinnamate produced from phenylalanine by PAL, while formation of volatile phenylpropanoid-related (C_6_-C_2_) compounds occurs directly from phenylalanine (Dudareva et al., 2013) (Figure 3D). Genes encoding enzymes for this metabolic network (Supplementary Table S5) are significantly enriched with up-regulated DEGs and with DMGs having increased H3K9ac (Figures 3D and 3E). Interestingly, increases in both H3K9ac levels and transcript levels occur at gene loci underlying the first few enzymatic steps in the pathway, notably phenylacetaldehyde synthase (PAAS), cinnamoyl-CoA ligase (CNL), cinnamoyl-CoA hydratase-dehydrogenase (CHD), and 3-ketoacyl-CoA thiolase (KAT). By contrast, the enzymes acting at the final steps of VOC formation producing methylbenzoate, benzylbenzoate, and phenylethylbenzoate are not induced at the transcript level (Figure 3D). These end-step enzymes may rely on post-translational regulation or simply on increased precursor flux to generate the high levels of their products (Clark et al., 2009; Dudareva et al., 2004). Therefore, compared to the primary metabolism pathway (Figure 3A) where nearly every enzymatic step is regulated at the chromatin level and transcript level, the regulation of the secondary metabolism pathway seems to show more target specificity with a preference of enzymes of the early steps of the pathway.

### Anthocyanin pathway

Certain floral VOCs such as eugenol and isoeugenol are synthesized from 4-coumaroyl-CoA, which is also utilized to produce pigment anthocyanin (Quattrocchio et al., 1999). In *P. hybrida* cv. Mitchell, the flowers emit high levels of VOCs but the petals lack pigmentation. Our examination of the anthocyanin biosynthesis pathway (Figure 4A; Supplementary Table S6) (Quattrocchio et al., 1993; Ferrer et al., 2008; Passeri et al., 2016; Bombarely et al., 2016) found that transcriptional repression shuts off the anthocyanin branch post-anthesis, possibly contributing to the channeling of carbon flux toward VOC production (Figure 4A). Indeed, anthocyanin pathway genes are significantly enriched with down-regulated DEGs (Figure 4C). At the chromatin level, increased H3K9ac deposition during corolla development is almost completely excluded from the anthocyanin synthesis pathway. Overall, these results demonstrate that during the maturation of petunia corolla, there is a corresponding deactivation of genes involved in anthocyanin production concurrent with the activation of VOC branches of the phenylpropanoid pathway.

**Figure 4.**
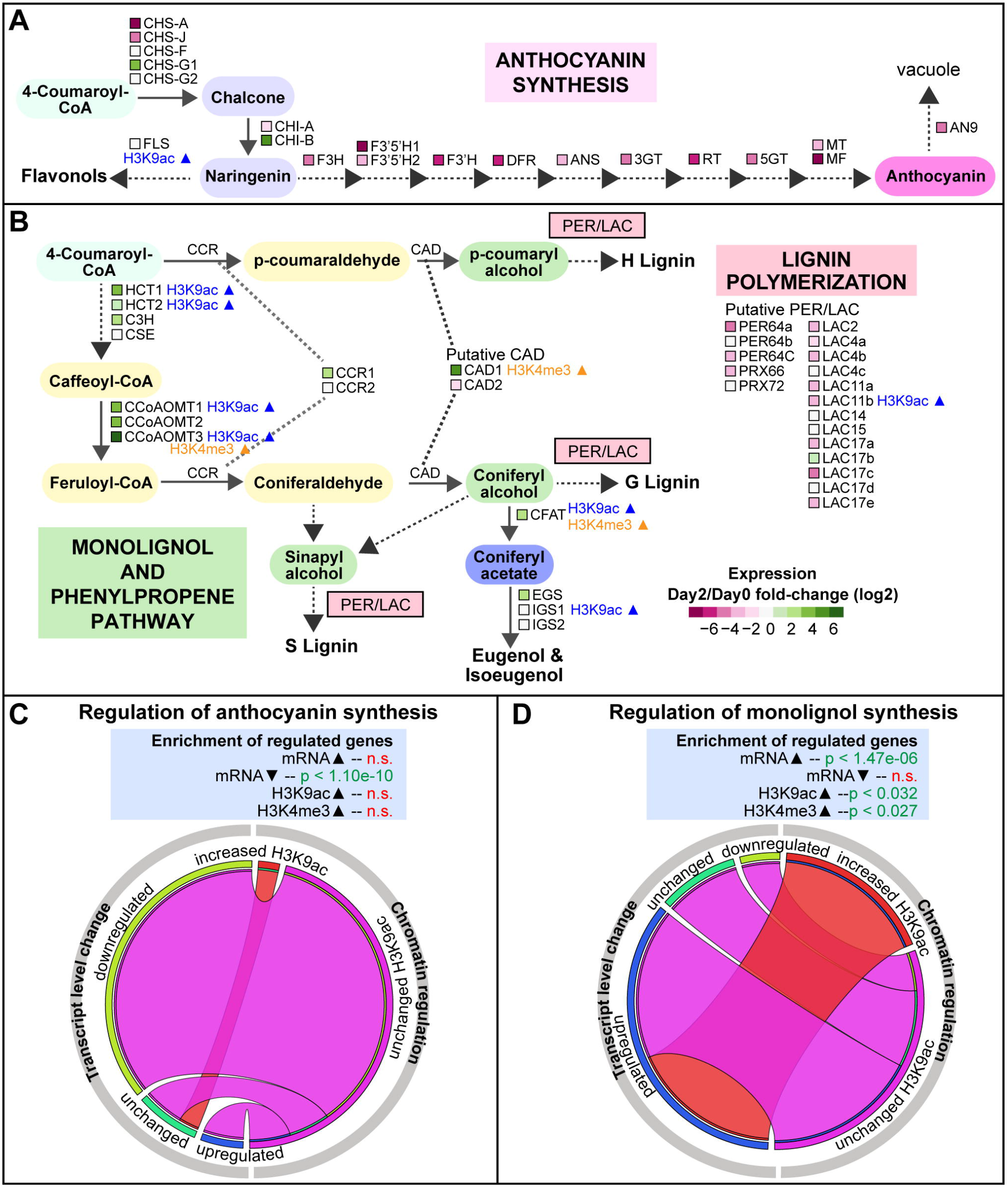
Transcriptional and epigenetic regulation ofanthocyanin, lignin, and eugenol/isoeugenol biosynthesis. **(A)** Schematic of the flavonoid and anthocyanin biosynthesis pathway with transcriptomic data overlaid. Color legend: general phenylpropanoid intermediate 4-coumaroyl-CoA in light blue, flavonoid branch intermediates in dark blue, anthocyanin in magenta. (B) Schematic of the monolignol pathway genes generating the three monolignol precursors, the eugenol and isoeugenol synthesis branch, and the lignin polymerization peroxidase/laccase genes, with transcriptomic and epigenomic data overlaid. Color legend: general phenylpropanoid intermediate 4-coumaroyl-CoA in light blue, monolignol branch intermediates in yellow, monolignol units in green, phenylpropene branch intermediate coniferyl acetate in dark blue. (C) Regulatory categories of anthocyanin pathway genes and hypergeometric p values determined for enrichment among DEGs and DMGs. (D) Regulatory categories of monolignol biosynthesis genes with hypergeometric p values determined for enrichment among DEGs and DMGs. 3GT, 3-glucosyl transferase; 5GT, 5-glucosyl transferase; ANS, anthocyanin synthase; C3H, coumarate 3-hydroxylase; CAD, cinnamyl alcohol dehydrogenase; CCoAOMT, caffeoyl-CoA O-methyltransferase; CCR, cinnamoyl-CoA reductase; CFAT, coniferyl alcohol acetyltransferase; CHI, chalcone isomerase; CHS, chalcone synthase; CSE, caffeoyl shikimate esterase; DFR, dihydroflavonol 4-reductase; EGS, eugenol synthase; F3’5’H, flavonoid 3’,5’-hydroxylase; F3H, flavanone 3-hydroxylase; F3’H, flavonoid 3’-hydroxylase; FLS, flavonol synthase; HCT, hydroxycinnamoyl-CoA:shikimate/quinate hydroxycinnamoyl transferase; IGS, isoeugenol synthase; LAC, laccase; MF, methylation at five; MT, methylation at three; PER/PRX, peroxidase; RT, rhamnosylation at three.

### Monolignol/phenylpropenes synthesis pathway

Our analysis revealed a complex and intriguing regulation pattern of the lignin biosynthetic branch of the phenylpropanoid pathway, with significant enrichment of lignin biosynthesis genes among both up-regulated and down-regulated genes on day 2 *vs* day 0 (Figure 2E). It should be noted, that volatile phenylpropene and lignin biosynthesis share the biochemical steps up to monolignol biosynthesis, where they diverge. As such, further investigation of the pathway revealed important regulatory control points to direct flux through the monolignol pathway to promote production of volatile eugenol and isoeugenol and repress synthesis of lignin polymers in the mature corolla.

In the monolignol biosynthetic pathway (Supplementary Table S7), enzymes including hydroxycinnamoyl-CoA:shikimate/quinate hydroxycinnamoyl transferase (HCT), caffeoyl-CoA O-methyltransferase (CCoAOMT), and coumarate 3-hydroxylase (C3H) utilize 4-coumaroyl-CoA from the general phenylpropanoid pathway to produce aldehyde intermediates p-coumaraldehyde or coniferaldehyde (Figure 4B) (Muhlemann et al., 2014b; Shaipulah et al., 2016; Kim et al., 2019). Cinnamyl alcohol dehydrogenase (CAD) enzyme(s) then convert the aldehyde intermediates to three monolignol intermediates, p-coumaryl alcohol, sinapyl alcohol, and coniferyl alcohol. Genes encoding these enzymes are induced on day 2 relative to day 0 (Figure 4B). Specifically, genes encoding the enzymes catalyzing the early steps (*HCT* and *CCoAOMT*) are targeted for H3K9ac deposition during corolla maturation (Figure 4B). Overall, there is a significant enrichment of up-regulated DEGs and DMGs with increased H3K9ac among the genes contributing to the production of the monolignol intermediates (Figure 4D), indicating that chromatin and transcriptional regulation underlie the increased flux through the monolignol pathway.

Two competing fates exist for these monolignol intermediates in petunia flowers. The monolignol can be used to synthesize lignin polymers for incorporation into the cell wall through the function of specialized peroxidases (PER) and laccases (LAC) (Fraser and Chapple, 2011) (Figure 4B). Alternatively, the monolignol coniferyl alcohol can serve as a precursor for production of the phenylpropene VOCs eugenol and isoeugenol (Muhlemann et al., 2014b), through the actions of coniferyl alcohol acetyltransferase (CFAT) (Dexter et al., 2007) and eugenol synthase (EGS) or isoeugenol synthase (IGS) (Koeduka et al., 2008) (Figure 4B). Both *CFAT* and *EGS* are induced in the corolla at the transcript level while *CFAT* and *IGS1* show increased H3K9ac levels. By contrast, we observed a clear pattern of decreased transcript levels for *PER* and *LAC* (Figures 4B, Supplementary Table S7), which is in agreement with low levels of lignification observed in petunia flower relative to stem and leaf tissue (Muhlemann et al., 2014b; Shaipulah et al., 2016).

Overall, our results suggest that important control points of the monolignol/phenylpropene metabolic network are regulated at the chromatin level, possibly to direct flux through the monolignol pathway to produce VOCs. The apparent GO term enrichment for lignin biosynthesis in both up- and down-regulated genes (Figure 2E) is due to the separate regulatory regimes governing the different modules of the lignin branch of phenylpropanoid metabolism; production of monolignol intermediates is activated to produce phenylpropene VOCs in petunia, while at the same time there is a decreased incorporation of monolignol subunits into lignin polymers.

### Histone acetylation is essential for transcriptional activation of VOC genes and VOC emission

To determine whether the observed dynamic histone acetylation has a causal effect on the induction of VOC genes, we treated petunia flower buds with chemical inhibitors of histone acetyltransferases (HATs) and tested whether the activation of VOC genes during anthesis was impeded. In detail, petunia buds on day 0 were treated with C646 or MB-3, which are small molecule inhibitors of HATs (Malapeira et al., 2012; Lee et al., 2015; Aquea et al., 2017). qRT-PCR was then performed to measure the gene expression on days 0, 1 and 2 for a group of genes with increased H3K9ac and transcriptional activation during anthesis in our ChIP-Seq and RNA-Seq analysis (Supplementary Datasets S1 and S2), selected from different branches of VOC pathways: phenylalanine biosynthesis (*EPSPS1*, *DAHP1*, *CS*), benzenoid VOC biosynthesis (*CNL1*, *KAT1*), and phenylpropene VOC biosynthesis (*CCoAOMT1*, *CFAT*). Also included were *ODORANT1 (ODO1)*, encoding a transcription factor involved in activating shikimate pathway and VOC genes (Verdonk et al., 2005), and *ATP-binding cassette subfamily G member 1* (*PhABCG1)*, encoding an active VOC transporter required for VOC emission (Adebesin et al., 2017). The tested genes showed activation of gene expression during anthesis in mock samples, and strikingly, all the genes exhibited a notable attenuation of transcriptional activation when treated with MB-3 (Figure 5A). The histone acetylation levels at select gene loci were measured at day 2 using ChIP-qPCR and H3K9ac showed significant reduction in the MB-3 treated samples compared to the mock treated samples, supporting that the attenuation of transcriptional activation was indeed due to reduced histone acetyltransferase activity (Figure 5B). H3K14ac levels were unaffected (Figure S4B), suggesting that MB-3 particularly affects the H3K9ac levels at this developmental stage. Together, these results support the mechanistic importance of H3K9ac deposition in activating primary and secondary metabolic pathways for VOC synthesis and emission in the petunia corolla. Treatment with the other inhibitor, C646, did not alter the transcriptional activation of any tested genes (Figure 5A). MB-3 inhibits the GNAT family HATs and to a lesser extent p300/CBP family HATs, while C646 mainly inhibits the p300/CBP family (Biel et al., 2004; Bowers et al., 2010). Our results thus suggest that unknown member(s) of the GNAT family likely have a specialized role in the transcriptional regulation of VOC biosynthesis.

**Figure 5.**
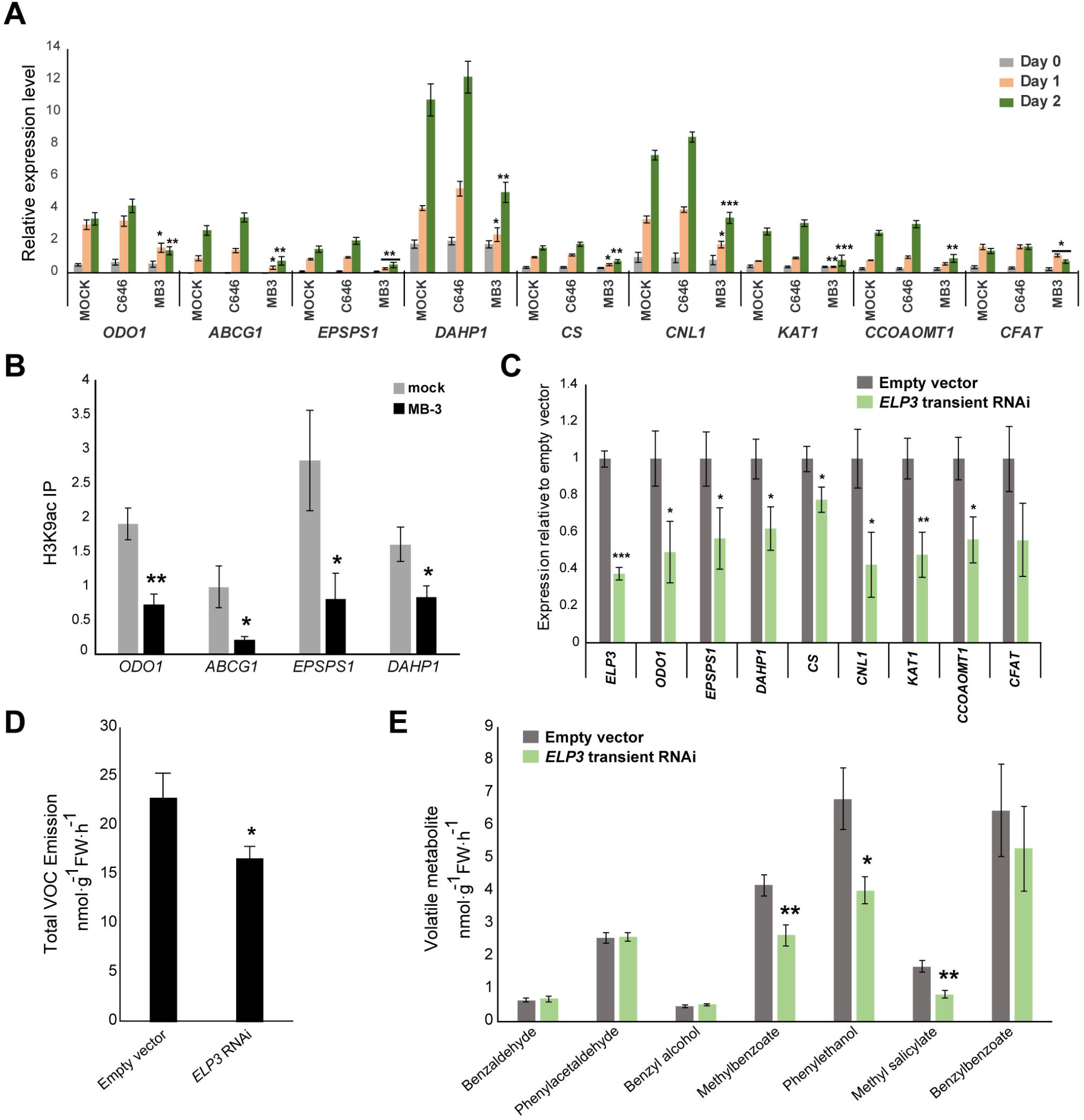
Inhibition ofhistone acetylation affects voe gene transcription and voe emission. (A) qRT-PeR results showing that the induction ofvoe genes is impaired by histone acetyltransferase inhibitor MB-3, but not by e646. The relative expression levels normalized to reference genes UBQl0 and FBP l are shown. Error bars represent standard error ofthe mean for biological replicates (n=3-4). (B) ehIP-qPeR results showing a significant reduction ofH3K9ac in the MB-3 treated samples compared to the mock samples. Detached flower buds at day 0 were treated with mock (0.1 % DMSO) or MB-3 (100 μM in 0.1 % DMSO) for two days and then processed for ehIP with antibodies against H3K9ac. qPeR was performed with primers designed for gene body around the acetylation peaks based on the ehIP-Seq data in this study. Immunoprecipitation (IP) ofehIP samples was first normalized to the input DNA and then normalized to housekeeping genes AeTIN and PP2AA3. Significance ofdifference between mock samples and MB-3 treated samples was determined by Student’s t-test (n=4) and depicted as asterisks: p < 0.05 (*), p < 0.01 (**). (e) qRT-PeR results showing that transient RNAi successfully suppresses the expression ofELP3 as well as reducing the expression level ofvoe genes. Six biological replicates ofindependent transformation events were included. The relative expression levels were determined by qRT-PeR, normalized to UBQIO and FBPl and scaled relative to empty vector (EV) control. Error bars represent standard error ofthe mean (n=6). (D and E) Floral voe emission from petunia flowers measured as total emission (D) or individual metabolites (E) in ELP3 transient RNAi samples and empty vector control. Significance ofdifference between treatment vs mock (A) or between ELP3 RNAi vs EV (e-E) is determined by Student’s t-test and depicted by asterisks: p ≤ 0.05 (*), p ≤ 0.01 (**), p ≤ 0.001 (***). Error bars represent standard error ofthe mean for biological replicates from independent transformations (n=6).

To determine which member of the GNAT family is responsible for the deposition of H3K9ac that influences VOC activation, the HAT family genes were identified in the *P. axillaris* genome based on homology with Arabidopsis HAT genes (Pandey et al., 2002) (Supplementary Table S8). Among the identified HAT genes, *ELP3*, a GNAT family HAT, was unique in being significantly up-regulated (FDR<0.05; fold change >2) during corolla development (Supplementary Dataset S2, Figure S5A). We thus hypothesized that ELP3 mediates VOC gene expression in petunia flower during anthesis. To test this hypothesis, petunia *ELP3* was down-regulated in flowers using transient RNA interference (RNAi) strategy. *ELP3* RNAi was performed in six biological replicates, each representing independent transformation events, and the expression of select VOC genes and levels of emitted VOCs were measured two days post-infiltration. We observed an average reduction by 62% in *ELP3* transcript levels (Figures 5C), with only minor changes in expression levels of other HAT genes (Figure S5B). Knockdown of *ELP3* resulted in significant down-regulation of most VOC biosynthetic genes assayed (Figure 5C), suggesting that the expression of these VOC genes is dependent on ELP3. Importantly, the total emitted volatiles were also decreased in *ELP3* RNAi flowers relative to empty vector control (Figure 5D), with individual compounds affected to different degrees (Figure 5E). Taken together, these RNAi results along with chemical inhibition experiments provide two lines of *in planta* evidence that histone acetylation, mediated at least in part by ELP3, activates transcription of VOC metabolic pathways, which is required for the synthesis and emission of volatile benzenoid products.

### Epigenetic and transcriptional activation of hormone signaling pathways during anthesis

To further investigate the regulatory and signaling mechanisms that govern VOC production and release, we examined GO terms involved in plant signaling for their significant enrichment among the DMGs and DEGs (Figure 6A). Interestingly, ‘positive regulation of gibberellic acid (GA) mediated signaling pathway’ is the only significantly enriched GO term shared between transcriptionally activated DEGs and DMGs gaining H3K9ac.

**Figure 6.**
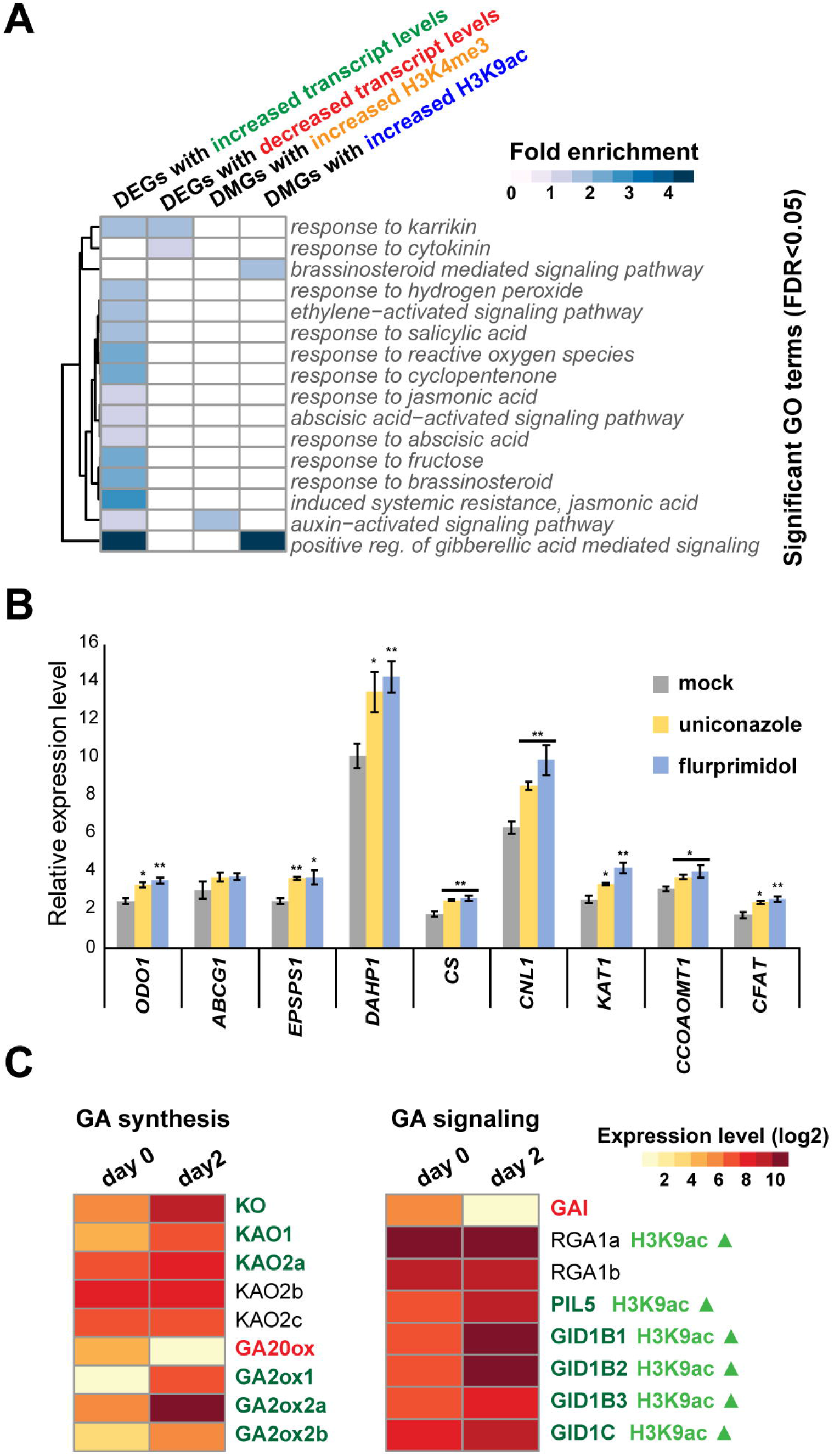
Phytohormone and Gibberellic Acid Signaling in Petunia Corolla. **(A)** Heatmap depicting the fold enrichment of signaling GO terms significantly enriched among the DEGs or DMGs. **(B)** qRT-PCR results showing the up-regulation ofVOC genes when GA synthesis is inhibited by uniconazole or flurprimidol. Relative expression levels normalized to UBQ10 and FBP1 are shown and error bars represent standard error of the mean for biological replicates (n=3-4). Significance of treatment relative to mock is determined by Student’s t-test and depicted as asterisks: p < 0.05 (*), p < 0.01 (**), p < 0.001 (***). (C) Heatmap depicting expression levels of GA synthesis and signaling genes determined by RNA-seq. Up-regulated DEGs are shown in green and down regulated DEGs are shown in red. Log2 normalized expression levels are shown for genes whose expression levels are above the first quartile of all genes in corolla at either day O or day 2. Genes names reflect best homology to Arabidopsis gene copies and multiple expressed homologs were named alphabetically a through c, or for GIDl B, copies 1 through 3. GAi, GA insensitive; GID, GA insensitive dwarf; KO, ent-kaurene oxidase; KAO, ent-kaurene acid oxidase; PIL, phytochrome-interacting factor 3-like; RGA, repressor of gal-3.

It has been reported that GA represses VOC synthesis and emission (Ravid et al., 2017). Thus, it would be reasonable to expect that GA biosynthesis and signaling are down-regulated when VOC emission is induced. In contrast, our results revealed up-regulation of GA signaling genes during the developmental timeframe of VOC induction (Figure 6A). To investigate this discrepancy, we first tested whether GA serves as a positive or negative regulator of VOC genes in our system. Flower buds were treated with 100 μM uniconazole or flurprimidol, inhibitors of GA synthesis, or mock control (Yamaguchi et al., 2001; Pullman et al., 2005), and the expression levels of selected VOC genes on day 2 were analyzed by qRT-PCR. Inhibition of GA biosynthesis led to a moderate but statistically significant increase in the transcript levels of most VOC genes assayed (Figure 6B). Therefore, our results agree with the previous report (Ravid et al. 2017) that GA represses VOC synthesis. Next, we took advantage of our genome-wide data to examine the temporal regulation of GA synthesis and GA signaling pathways during anthesis (Figure 6C, Supplementary Table S9). Interestingly, GA biosynthesis and GA signaling genes show distinct regulation patterns. The GA biosynthetic genes show mixed expression patterns and lack dynamic histone acetylation (Figure 6C). Notably, although *ent-KAURENE OXIDASE* (*KO*) and *ent-KAURENE ACID OXIDASE* (*KAO*) are well expressed, *GA20ox*, which acts downstream of KAO to produce bioactive GA, is strongly down-regulated from day 0 to day 2 (Figure 6C). Meanwhile, several copies of *GA2ox*, which convert bioactive GA to inactive forms, are significantly and strongly induced from day 0 to day 2 (Figure 6C). Our data thus suggest that the biosynthesis of GA in corolla is down-regulated post-anthesis. In contrast, many genes involved in GA perception and signaling are up-regulated with increased H3K9ac during anthesis, notably including four copies of GA receptor *GIBBERELLIN INSENSITIVE DWARF1 (GID1)* (Tyler et al., 2004; Murase et al., 2008), one copy of *REPRESSOR OF GA 1-3* (*RGA1)*, and *PHYTOCHROME INTERACTING FACTOR3-LIKE 5 (PIL5)*, a positive regulator of *RGA1* (Oh et al., 2007, 5). Overall, our integrated ChIP-Seq and RNA-Seq analyses uncovered an intriguing role of GA signaling in regulating VOC production, suggesting low GA synthesis but high GA sensitivity during peak VOC emission. It is possible that GA may repress the VOC pathway before the flower opens, and the reduction in GA levels after flower opening allows high levels of VOC emission. Meanwhile, the corolla tissues become highly sensitive to GA, as the GA level may rise again during senescence or after pollination to shut off the VOC emission.

## Discussion

Rapid and efficient production of secondary metabolites in response to developmental and environmental stimuli requires coordinated regulation of primary and secondary metabolic pathways. By integrating ChIP-Seq and RNA-Seq analyses, we have shown that chromatin level regulation acts as an underlying mechanism activating primary and secondary metabolic networks leading to formation of floral VOCs during flower development in petunia. Our data revealed an exquisite coordination balancing substrate availability and flux via different metabolite fates through dynamic regulation of chromatin modification and transcript levels of metabolic gene networks (Figure 7). While chromatin modification and transcriptional reprogramming ensures both that large amounts of phenylalanine substrate are available and that it flows preferentially to VOC biosynthesis, other metabolic fates are blocked by transcriptional inactivation of those branches. In detail, during flower opening and post-anthesis, H3K9ac is deposited at gene loci encoding enzymes involved in the shikimate and phenylalanine biosynthesis pathways, with a corresponding increase in transcriptional activity, enabling the production of large amounts of phenylalanine (Figures 3A and 7). Phenylalanine is in turn a precursor for the volatile benzenoid and phenylpropanoid network, the enzymes involved in which are regulated in a targeted manner through increased H3K9ac and up-regulated transcript level to direct metabolic flux to VOC production (Figures 3D and 7). Phenylalanine is also converted to 4-coumaroyl-CoA, which is a common substrate of divergent phenylpropanoid branches including flavonoid/anthocyanin synthesis and monolignol biosynthesis. Targeted H3K9ac and transcript level increases are observed in the monolignol synthesis pathway and the enzymes that produce the phenylpropene VOCs eugenol and isoeugenol (Figures 4B and 7), while there is a simultaneous deactivation of lignin polymerization (Figures 4B and 7) and anthocyanin biosynthesis (Figures 4A and 7), with the net effect of directing 4-coumaroyl-CoA substrate toward phenylpropene VOC production.

**Figure 7.**
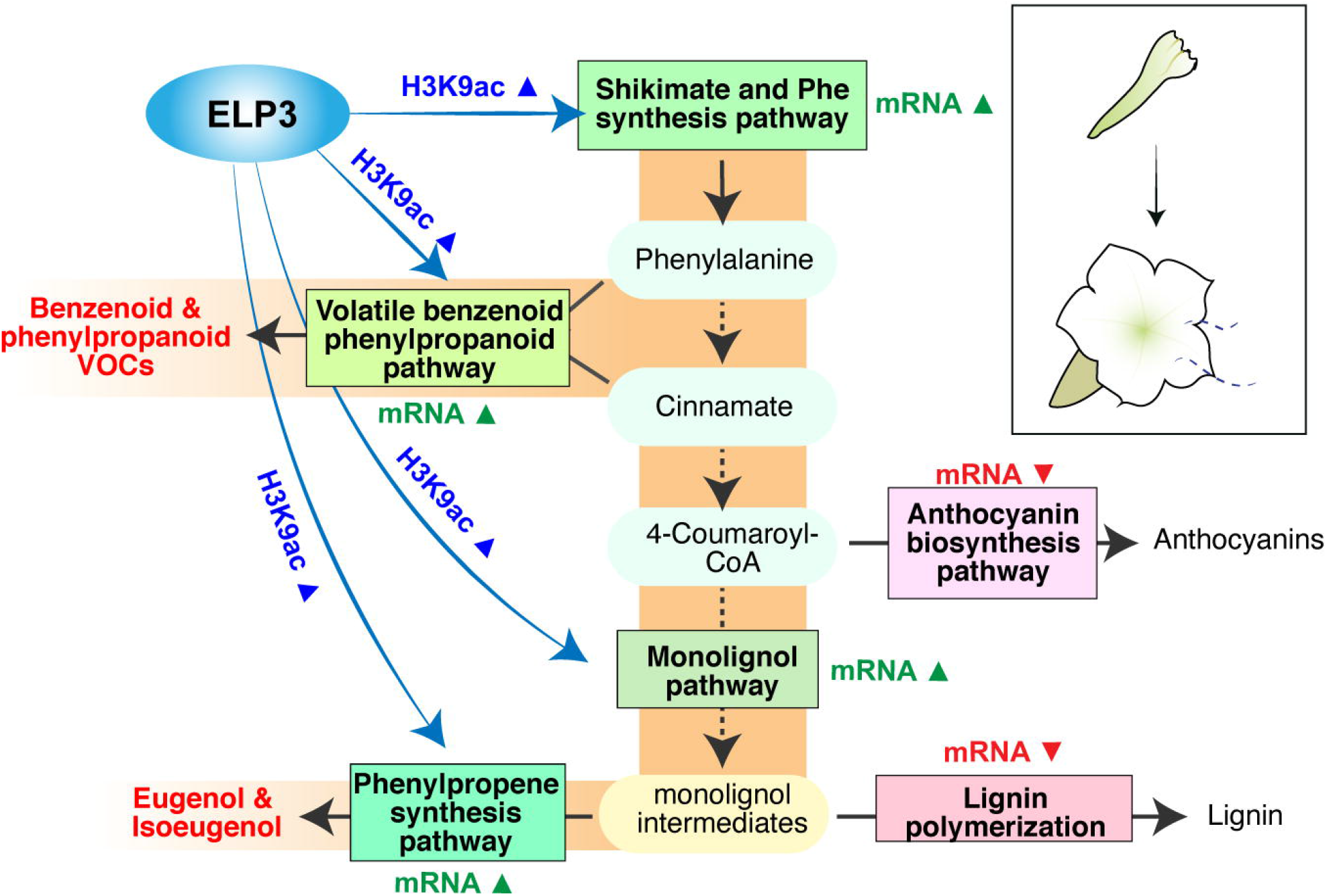
Summary Model of Chromatin Level Regulation and Transcriptional Reprogramming to Facilitate VOC Emission During Petunia Corolla Development. This schematic summarizes regulation of phenylalanine and phenylpropanoid synthesis branches observed at the chromatin level through increased histone acetylation (H3K9ac) at day 2 post-anthesis compared to day O bud tissue in the petunia corolla, and corresponding transcriptional changes in order to direct metabolic flow through the phenylpropanoid pathway to generate floral VOCs.

Our findings in petunia share similarity with the epigenetic control of secondary metabolism in fungi, where histone acetyltransferases contribute to the transcriptional activation of biosynthetic gene clusters and production of a range of specific secondary metabolites (Strauss and Reyes-Dominguez, 2011; Pfannenstiel and Keller, 2019). Our data supports that H3K9ac is a specific histone mark contributing toward activating the VOC synthesis pathway, rather than merely a permissive histone mark associated with activated genes. Importantly, blocking HAT activity by chemical inhibition attenuates the normal transcriptional induction of these pathways, thus supporting that *de novo* histone acetylation is required for the VOC synthesis (Figure 5). Indeed, another common permissive histone mark assayed in our study, H3K4me3, was not significantly associated with VOC production during anthesis. One possible explanation for this specific role of histone acetylation is that among the different histone modifications, histone acetylation is hypothesized to integrate information on cellular metabolic status into chromatin states because it uses acetyl-coA as substrate (Dutta et al., 2016; Reid et al., 2017).

Histone acetylation is “written” by HAT family proteins. We found that a specific HAT, ELP3, activates multiple genes in the VOC gene network (Figure 7), by showing that transient RNAi knockdown of *ELP3* attenuates the transcription of genes in the VOC pathways as well as the level of emitted volatile metabolites (Figures 5C-E). ELP3, short for Elongator complex subunit 3, was first discovered through association with RNAPII in yeast (Otero et al., 1999; Krogan and Greenblatt, 2001), and later was found to be conserved among archaea, bacteria, yeast, animals and plants. The elongator complex consists of six subunits, of which ELP3 is the catalytic subunit with HAT activity. The *elp3* mutants in plants have narrow leaves (Nelissen et al., 2005; Nelissen et al., 2010) and altered root morphology (Jia et al., 2015; Qi et al., 2019). Previous studies suggest a role of ELP3 in regulating gene transcription with target specificity primarily in dividing meristemic tissues (Nelissen et al., 2010; Jia et al., 2015; Qi et al., 2019). Our findings now link ELP3 with activation of the VOC metabolic network in mature flowers, where it is likely responsible for deposition of H3K9ac along the gene body of VOC genes to facilitate high levels of transcription.

The mechanistic insights obtained in this study on petunia corolla provide a framework to understand epigenetic regulatory mechanisms underlying plant secondary metabolic pathways, the activity of which are often restricted by developmental and environmental contexts. Importantly, when VOC pathways are activated in the corolla, coordinated H3K9ac deposition and transcriptional activation mainly occurs at gene loci encoding enzymes critical for directing flow of shared precursor metabolites toward VOC formation. This is the case for *DAHP1* in the shikimate pathway (Figure 3A), *CM1* in phenylalanine biosynthesis (Figure 3A), *PAAS1* and *CNL1* in volatile benzenoid and phenylpropanoid biosynthesis (Figure 3D), *HCT1* and *HCT2* in monolignol biosynthesis (Figure 4B), and *CFAT* in eugenol and isoeugenol biosynthesis (Figure 4B). How such target specificity is achieved, from biochemical to evolutionary scales, is an intriguing question that requires further investigation.

Finally, the uncovered chromatin level regulation of secondary metabolic pathways has practical implications in metabolic engineering. Engineering of phenylpropanoid and VOC metabolism has been a topic of interest (Brilli et al., 2019; Dudareva and Negre, 2005; Plasmeijer et al., 2020), as VOCs provide economically important flavor and scent characteristics to food and botanical products (Dudareva et al., 2013). It was noted that an explicit control of metabolic flow through targeted primary and secondary metabolic pathways is required in order to produce high levels of desired specialized metabolites. Our results showed that these metabolic networks are regulated at chromatin level through histone modifications such as H3K9ac. Therefore, the optimal outcomes of metabolic engineering will require a mindset toward network level regulation of primary and secondary metabolism, and efforts may be critically impacted by the local chromatin environment of transgenes.

## ABBREVIATIONS

HAT: histone acetyltransferase
VOC: volatile organic compound
H3K9ac: Histone subunit 3 lysine 9 acetylation
H3K4me3: Histone subunit 3 lysine 4 trimethylation
GNAT/MYST: Gcn5-related N-acetyltransferase/ MOZ, Ybf2/Sas3, Sas2, Tip60 family
ELP3: Elongator complex subunit 3
RNAi: RNA interference
ChIP-Seq: Chromatin Immunoprecipitation Sequencing
GO: Gene Ontology
FDR: False discovery rate
GC-MS: Gas chromatography–mass spectrometry
DMGs: differentially modified genes
DEGs: differentially expressed genes

## Supplementary data

Supplementary data are available at JXB online.

Supplementary Figure S1. ChIP-Seq reads alignment to petunia genomes

Supplementary Figure S2. RNA-Seq reads alignment to petunia genomes

Supplementary Figure S3. Decreased H3K9ac DMGs

Supplementary Figure S4. ChIP-qPCR of MB-3 treated petunia

Supplementary Figure S5. Expression of histone acetyltransferases (HATs) in petunia

Supplementary Dataset S1. Increased and decreased H3K9ac and H3K4me3 DMGs

Supplementary Dataset S2. Transcriptionally activated and repressed DEGs

Supplementary Table S1. Primers used in the study.

Supplementary Table S2. Enriched GO Terms Among DMGs with increased H3K9ac.

Supplementary Table S3. Enriched GO Terms among DMGs with increased H3K4me3

Supplementary Table S4. Enriched GO terms among up-regulated (A) or down-regulated (B) DEGs.

Supplementary Table S5. Genes involved in shikimate and phenylalanine synthesis (a), general phenylpropanoid pathway (b) or volatile benzenoid and phenylpropanoid synthesis genes (c).

Supplementary Table S6. Flavonoid and anthocyanin synthesis genes

Supplementary Table S7. Genes involved in monolignol pathway, eugenol/isoeugenol synthesis, or lignin polymerization.

Supplementary Table S8. Histone acetyltransferase genes

Supplementary Table S9. Gibberellic acid signaling and metabolism genes

## Data Availability

ChIP-Seq data generated in this work are available at the NCBI Sequence Read Archive under BioProject accession PRJNA650505.

## Acknowledgments

This research is supported by a seed grant to Y.L. and N.D. from the Purdue Center for Plant Biology, USDA National Institute of Food and Agriculture Hatch project numbers 1013620 to Y.L., 177845 to N.D., and an Agriculture and Food Research Initiative Postdoctoral Fellowship (grant no. 2019-67012-29660) to R.M.P. from the USDA National Institute of Food and Agriculture. We thank Akhila Rose Prabin Kingsly for her technical assistance to grow plants.

## Author Contributions

R.M.P., Y.L., and N.D. conceived and designed the study. R.M.P. performed the experiments and data analysis. X.Q.H. performed transient RNAi experiments and metabolite profiling. R.M.P, X.Q.H., Y.L., and N.D. wrote and revised the manuscript.

